# Evaluation of self-generated behavior: untangling metacognitive read-out and error detection

**DOI:** 10.1101/513242

**Authors:** Tadeusz W. Kononowicz, Virginie van Wassenhove

## Abstract

When producing a duration, *for instance* by pressing a key for one second, the brain relies on self-generated neuronal dynamics to monitor the “*flow of time*”. Converging evidence has suggested that the brain can also monitor itself monitoring time. Here, we investigated which brain mechanisms support metacognitive inferences when self-generating timing behavior. Although studies have shown that participants can reliably detect temporal errors when generating a duration (Akdogan & Balci, 2017; Kononowicz et al., 2017), the neural bases underlying the evaluation and the monitoring of this self-generated temporal behavior are unknown. Theories of psychological time have also remained silent about such self-evaluation abilities. How are temporal errors inferred on the basis of purely internally driven brain dynamics without external reference for time? We contrasted the *error-detection* hypothesis, in which error-detection would result from the comparison of competing motor plans with the *read-out* hypothesis, in which errors would result from inferring the state of an internal code for motor timing. Human participants generated a time interval, and evaluated the magnitude of their timing (first and second order behavioral judgments, respectively) while being recorded with time-resolved neuroimaging. Focusing on the neural signatures following the termination of self-generated duration, we found several regions involved in performance monitoring, which displayed a linear association between the power of α (8-14 Hz) oscillations, and the duration of the produced interval. Altogether, our results support the read-out hypothesis and indicate that first-order signals may be integrated for the evaluation of self-generated behavior.

**SIGNIFICANCE STATEMENT:** When typing on a keyboard, the brain estimates *where* the finger should land, but also *when*. The endogenous generation of the *when* in time is naturally accompanied by timing errors which, quite remarkably, participants can accurately rate as being too short or too long, and also by how much. Here, we explored the brain mechanisms supporting such temporal metacognitive inferences. For this, we contrasted two working hypotheses (*error-detection vs. read-out*), and showed that the pattern of evoked and oscillatory brain activity parsimoniously accounted best for a read-out mechanism. Our results suggest the existence of meta-representations of time estimates.

## INTRODUCTION

Metacognition refers to the knowledge gained in introspecting about one’s cognitive states (Flavell 1979; Fleming & Dolan, 2012). Metacognition is often investigated through the evaluation of confidence on a perceptual decision task (*e.g.* discrimination of stimuli) thereby a second-order decision is contingent on a first-order judgment. Metacognition thus constitutes a meta-representation of the first-order judgement (Fleming, Dolan, Frith, 2012). Here, we explored the meta-representation of endogenous timing. In a seminal study, human participants receiving incorrect feedback following time production showed a negative evoked brain response post-feedback (Miltner *et al.*, 1997; then coined error-related negativity (ERN), now corresponding to feedback-related negativity). The ERN was interpreted as reflecting the difference between participants’ internal belief about the correctness of their time production, and the objective feedback. These observations suggested the possibility that an internal variable coding for duration could be studied from the perspective of metacognition but existing theories of psychological time have remained silent about the possibility of introspecting or self-evaluating internal representations of time.

Since, empirical evidence across species have converged on the possibility that time estimates were available for self-evaluation: for instance, the combination of uncertainty estimates about exogenous and endogenous timing may serve temporal monitoring (Balci et al, 2009). Metacognitive abilities in rats showed that, in a duration discrimination task, individuals more readily declined the test when provided with uncertain stimuli (Foote & Crystal, 2007). Additionally, humans can reliably report their temporal errors following time reproduction (*i.e.*, motor reproduction of a sensory time interval; Akdogan & Balci, 2017) and production (*i.e.*, self-generation of a time interval in the absence of sensory template, Fig. 1a; Kononowicz et al., 2017). In the latter, the precision of self-generated time intervals (first-order judgment, FOJ) informed participants’ self-evaluation (second-order judgment, SOJ) yielding accurate estimates of the signed magnitude of temporal errors.

**Figure 1.**
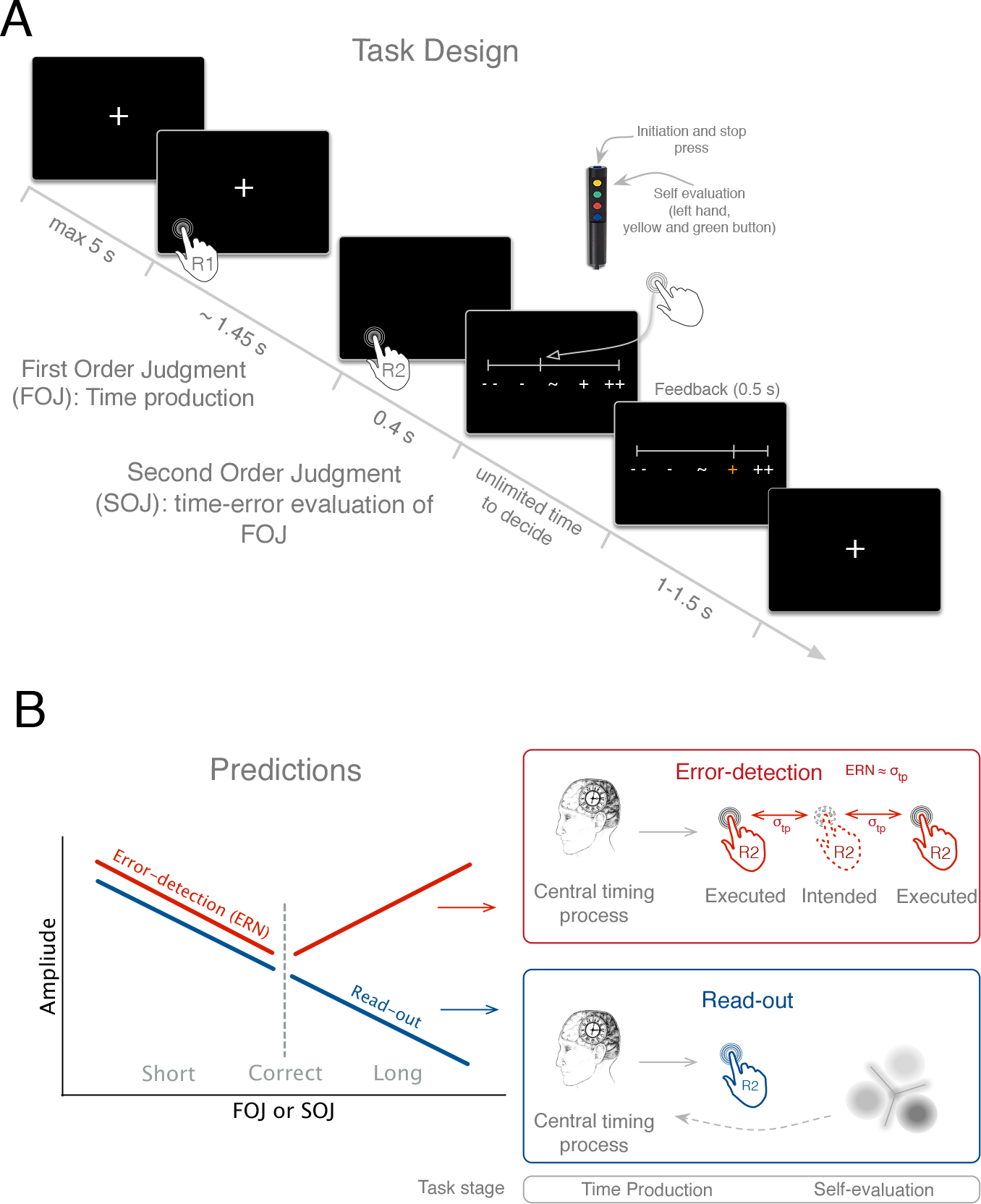
Experimental paradigm and predictions. (**A**) Time course of an experimental trial. (**B**) Predictions of the *temporal error-detection* (TED; red) and *temporal metacognitive read-out* (TMC; blue) working hypotheses. According to TED, temporal error evaluation arises from the online comparison of the planned and the actualized action in the temporal dimension. The dashed hand depicts intended action. The solid line hands depict misalignment of executed action with respect to the intendent one. As stated in the figure ERN amplitude would be proportional to the size of this misalignment between executed and intended actions (ERN ≈ σ_tp_). According to TMC, temporal error evaluation elicits a metarepresentation coding for a target duration by inferring the state of the networks that code for produced duration in the first place. Contrary to the TED, which was expected to be linked with ERN amplitude modulations, TMC was expected to elicit more sustained components as power modulations. Slower components would suggest involvement of processes other than temporal error detection. A direct read-out of internal variable would predict a linear scaling between the produced duration and the neural responses elicited after the temporal production (blue).

To date, the mechanisms supporting the evaluation of temporal error is unknown. Here, we investigated the neural mechanisms underlying temporal evaluation and contrasted two working hypotheses: (i) *temporal error-detection* of motor plans (Meckler et al., 2010; Praamstra et al., 2003) and (ii) *read-out* of an internal variable coding for duration (Fig. 2). The *temporal error-detection (TED)* hypothesis entails the comparison between the intended and the executed action: *TED* would occur following the end of the temporal production (R2), and was predicted to elicit the ERN (Cohen, 2014; Gehring et al., 1993). Late motor responses in deadline reaction time tasks elicit larger ERNs than early responses, suggesting a real-time monitoring of the unfolding action (Luu et al., 2010). During sensorimotor synchronization, the amplitude of the ERN increases for errors irrespective of their being early or late (Jantzen et al., 2017). Under the TED hypothesis, a V-shaped pattern was predicted so that the further away time productions were from the target, the larger the amplitude of the ERN (Fig. 1b). Under the TED hypothesis, the ERN would increase with motor errors and result from the real-time monitoring of action plans. To the contrary, *temporal metacognition* (*TMC*) assumes a meta-representation that actively infers the state of an internal variable for a target duration: a direct read-out of this internal variable would predict a linear scaling between the produced duration and the neural responses elicited after the temporal production (post-R2). In other words, a readout was predicted to code for the state of the networks related to duration estimation at the outset of the timed interval (Ivry & Schlerf, 2008; Karmarkar & Buonomano, 2007; Laje & Buonomano, 2013; Simen et al., 2011).

**Figure 2.**
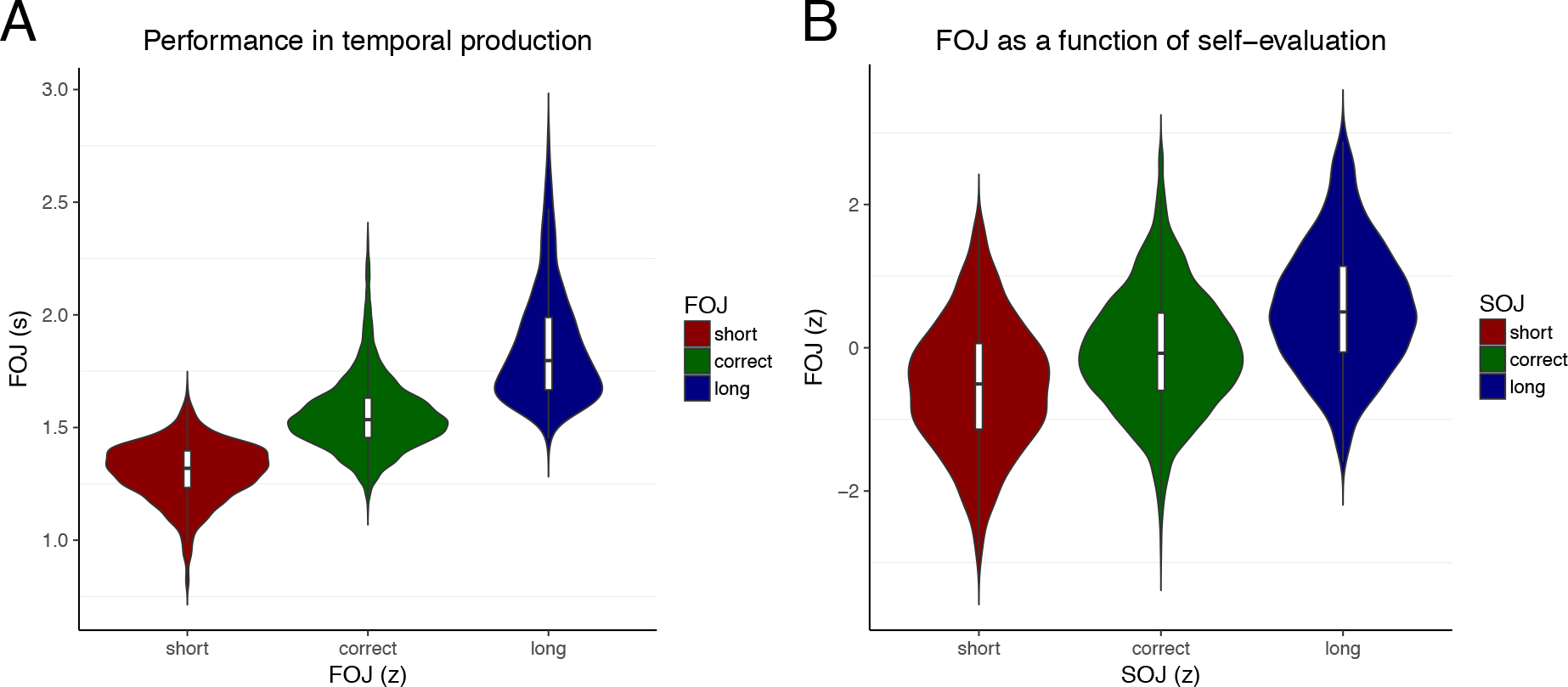
Temporal productions (FOJ) and self-evaluations (SOJ). **(A)** Participants accurately produced the target duration. The FOJ (first-order judgements) were clustered on the basis of the normalized FOJ yielding the ‘short’, ‘correct’ and ‘long’ categories. **(B)** FOJ plotted as a function of SOJ show a clear increase indicating that participants accurately self-evaluated (SOJ) their produced durations (FOJ).

We provide behavioral evidence demonstrating that participants accurately tracked their timing performance on a trial-by-trial basis during a time production task. We quantified brain evoked and oscillatory responses and found a novel link between neural markers of timing-related decision processes (*i.e.*, slow evoked activity and β (15-40 Hz) activity) during time production (FOJ) and metacognitive processes (SOJ).

## METHODS

### Participants

Nineteen right-handed volunteers (11 females, mean age: 24 years old) with no self-reported hearing/vision loss or neurological pathology were recruited for the experiment and received monetary compensation for their participation. Prior to the experiment, each participant provided a written informed consent in accordance with the Declaration of Helsinki (2008) and the Ethics Committee on Human Research at Neurospin (Gif-sur-Yvette). The data of seven participants were excluded from the analysis due to the absence of anatomical MRI, technical issues with the head positioning system during MEG acquisition, abnormal artifacts during MEG recordings, and two participants not finishing the experiment. These datasets were excluded *a priori*, and were neither visualized nor inspected. The final sample comprised twelve participants (7 females, mean age: 24 y.o.). All participants performed six experimental blocks, two participants performed five experimental blocks.

### Stimuli and Procedure

Before the MEG acquisitions, participants were explained they were taking part in a time estimation experiment, and written instructions were provided explaining all steps of the experimental protocol. In each trial, participants produced a 1.45 s time interval, rated on a linear scale whether their production was too short or too long as compared to the target interval, and finally, received feedback on their time production (not on their self-estimation) (Fig. 1a). We will refer to the produced time interval as the first order temporal judgment (FOJ), and to the self-estimation of the first order judgment as the second order temporal judgment (SOJ).

Participants received feeback on all trials in the 1^st^ and 4^th^ experimental blocks, and on 15% of the trials in all other blocks (Fig. 1a). To tailor an accurate feedback for each individual, a perceptual threshold for duration discrimination of the same 1.45 s duration was collected before the experiment. The individualized thresholds were used to scale the spacing of feedback categories for too short, correct or too long (Fig. 1b) and, unbeknownst to the participants, to adjust feedback in blocks 4 to 6.

Each trial started with the presentation of a fixation cross “+” on the screen indicating participants they could start whenever they decided to (Fig. 1a). The inter-trial interval ranged between 1 s and 1.5 s. Participants initiated their time production with a brief but strong button press once they felt relaxed and ready to start. Once they estimated that a 1.45 s interval had elapsed, they terminated the interval by another brief button press. To initiate and terminate their time production (FOJ) participants were asked to press the top button of a Fiber Optic Response Pad (FORP, Science Plus Group, DE) using their right thumb (Fig. 1b). The “+” was removed from the screen during the estimation of the time interval to avoid any sensory cue or confounding responses in brain activity related to the FOJ.

Following the production of the time interval, participants were asked to self-estimate their time estimation (second order judgment; Fig 1b). For this, participants were provided with a scale displayed on the screen 0.4 s after the keypress that terminated the produced time interval. Participants could move a cursor continuously using the yellow and green FORP buttons (Fig. 1b). Participants were instructed to place the cursor according to how close they thought their FOJ was with respect to the instructed target interval indicated by the sign ‘~’ placed in the middle of the scale. Participants placed the cursor to indicate whether they considered their produced time interval to be too short (‘−−’, left side of the scale) or too long (‘++’, right side of the scale). Participants could take as much time as needed to be accurate in their SOJ.

Following the completion of the SOJ, participants received feedback displayed on a scale identical to the one used for SOJ. The row of five symbols indicated the length of the just produced FOJ (Fig. 1a). The feedback range was set to the value of the perceptual threshold estimated on a per individual basis (mean population threshold = 0.223 s, SD = 0.111 s). A near correct FOJ yielded the middle ‘~’ symbol to turn green; a too short or too long FOJ turned the symbols ‘−’ or ‘+’ orange, respectively (Fig. 1b); a FOJ that exceeded these categories turned the symbols ‘−−’ or ‘++’ red. In Block 1 and 4, participants received feedback in all trials; in Block 2, 3, 5 and 6, participants received feedback in 15% of randomly selected trials (Fig. 1a). From Block 4 on, and unbeknownst to participants, the target duration was increased to 1.45 s + *individual threshold*/2 (mean population duration = 1.56 s). In Block 1 and 4, participants had to produce 100 trials; in Block 2, 3, 5, and 6, participants produced 118 trials. Between the experimental blocks, participants were reminded to produce the target duration of 1.45 s as accurately as possible, and to maximize the number of correct trials in each block.

### Estimation of temporal discrimination thresholds

The psychoacoustics toolbox was used to calculate the temporal discrimination threshold for each participant (Soranzo & Grassi, 2014) by adapting the available routine “DurationDiscriminationPureTone” provided in the toolbox. An adaptive procedure was chosen using a staircase method with a two-down one-up rule, and stopped after twelve reversals (Levitt, 1971). For each trial, three identical tones of 1 kHz were presented to the participants. One of the tones lasted longer than 1.45 sec (deviant tone) while the other two tones lasted precisely 1.45 sec (standard tones). The position of the deviant tone changed randomly across trials. The task was to identify the deviant tone and to give its position in the sequence. Tones were provided by earphones binaurally. The value of the correct category was set as *target duration* +/− (*threshold*/3), and the lower and upper limits were set as *target duration* +/− (2* *individual threshold*/3), respectively. These values were used to provide feedback to participants.

While this method did not provide a direct assessment of an individual’s temporal production discrimination threshold, the link between auditory and motor timing has been noted (e.g., Meegan, Aslin, Jacobs, 2000) and functionally relevant (*e.g.* Zatorre, Chen, Penhune, 2007).

### Simultaneous M/EEG recordings

The experiment was conducted in a dimly-lit, standard magnetically-shielded room located at Neurospin (CEA/DRF) in Gif-sur-Yvette. Participants sat in an armchair with eyes open looking at a screen used to show visual stimuli using a projector located outside of the magnetically shielded room. Participants were asked to respond by pushing a button on a FORP response pad (Science Plus Group, DE) held in their right hand. Electromagnetic brain activity was recorded using the whole-head Elekta Neuromag Vector View 306 MEG system (Neuromag Elekta LTD, Helsinki) equipped with 102 triple-sensors elements (two orthogonal planar gradiometers, and one magnetometer per sensor location) and the 64 native EEG system using Ag-AgCl electrodes (EasyCap, Germany) with impedances below 15 kΩ. Participants sat in upright position. Their head position in the dewar was measured before each block using four head-position coils placed over the frontal and the mastoid areas. The four head-position coils and three additional fiducial points (nasion, left and right pre-auricular areas) were digitized for subsequent co-registration with the individual’s anatomical MRI. MEG and EEG (M/EEG) recordings were sampled at 1 kHz and band-pass filtered between 0.03 Hz and 330 Hz. The electro-occulograms (EOG, horizontal and vertical eye movements), -cardiograms (ECG), and -myograms (EMG) were recorded simultaneously with MEG. The head position with respect to the MEG sensors was measured using coils attached to the scalp. The locations of the coils and EEG electrodes were digitized with respect to three anatomical landmarks using a 3D digitizer (Polhemus, US/Canada). Stimuli were presented using a PC running Psychtoolbox software (Brainard, 1997) that has been executed in Matlab environment.

## Data Analysis

### M/EEG data preprocessing

Signal Space Separation (SSS) correction (Taulu & Simola, 2006), head movement compensation, and bad channel rejection was done using MaxFilter Software (Elekta Neuromag). Trials containing excessive ocular artifacts, movement artifacts, amplifier saturation, or SQUID artifacts were automatically rejected using rejection criterion applied on magnetometers (55e^−12^ T/m) and on EEG channels (250e^−6^ V). Trial rejection was performed using epochs ranging from − 0.8 s to 2.5 s following the first press initiating the time production trial. Eye blinks, heart beats, and muscle artifacts were corrected using Independent Component Analysis (Bell & Sejnowski, 1995) with mne-python. Baseline correction was applied using the mean value ranging from −0.3 s to −0.1 s before the first key press.

Preprocessed M/EEG data were analyzed using MNE Python 0.13 (Gramfort et al., 2014) and custom written Python code. For the analysis of evoked responses in the time domain, a low-pass zero phase lag FIR filter (40 Hz) was applied to raw M/EEG data. For time-frequency analyses, raw data were filtered using a double-pass bandpass FIR filter (0.8 – 160 Hz). The high-pass cutoff was added to remove slow trends, which could lead to instabilities in time-frequency analyses. To reduce the dimensionality, all evoked and time-frequency analyses were performed on virtual sensor data combining magnetometers and gradiometers into single MEG sensor types using ‘*as_type’* method from MNE-Python 0.13 for gradiometers. This procedure largely simplified visualization and statistical analysis without losing information provided by all types of MEG sensors (gradiometers and magnetometers).

### M/EEG-aMRI coregistration

Anatomical Magnetic Resonance Imaging (aMRI) was used to provide high-resolution structural images of each individual’s brain. The anatomical MRI was recorded using a 3-T Siemens Trio MRI scanner. Parameters of the sequence were: voxel size: 1.0 × 1.0 × 1.1 mm; acquisition time: 466s; repetition time TR = 2300 ms; and echo time TE= 2.98 ms. Volumetric segmentation of participants’ anatomical MRI and cortical surface reconstruction was performed with the FreeSurfer software (http://surfer.nmr.mgh.harvard.edu/). A multi-echo FLASH pulse sequence with two flip angles (5 and 30 degrees) was also acquired (Jovicich et al., 2006; Fischl et al., 2004) to improve co-registration between EEG and aMRI. These procedures were used for group analysis with the MNE suite software (Gramfort et al., 2014). The co-registration of the M/EEG data with the individual’s structural MRI was carried out by realigning the digitized fiducial points with MRI slices. Using mne_analyze within the MNE suite, digitized fiducial points were aligned manually with the multimodal markers on the automatically extracted scalp of the participant. To insure reliable coregistration, an iterative refinement procedure was used to realign all digitized points with the individual’s scalp.

### MEG source reconstruction

Individual forward solutions for all source locations located on the cortical sheet were computed using a 3-layers boundary element model (BEM) constrained by the individual’s aMRI. Cortical surfaces extracted with FreeSurfer were sub-sampled to 10,242 equally spaced sources on each hemisphere (3.1 mm between sources). The noise covariance matrix for each individual was estimated from the baseline activity of all trials and all conditions. The forward solution, the noise covariance and source covariance matrices were used to calculate the dSPM estimates (Dale et al., 2000). The inverse computation was done using a loose orientation constraint (loose = 0.4, depth = 0.8) on the radial component of the signal. Individuals’ current source estimates were registered on the Freesurfer average brain for surface based analysis and visualization.

### ERF/P analysis

The analyses of MEG evoked-related fields (ERF) and EEG potentials (ERP) focused on the quantification of the amplitude of slow evoked components using non-parametric cluster-based permutation tests which control for multiple comparisons (Maris & Oostenveld, 2007. This analysis combined all sensors and electrodes into the analysis without predefining a particular subset of electrodes or sensors, thus keeping the set of MEG and EEG data as similar and consistent as possible. We used a period ranging from − 0.3 s to − 0.1 s before the first press as the baseline.

### Time-frequency analysis

To analyze the oscillatory power in different frequency bands using cluster based permutation, we used DPSS tapers with an adaptive time window of *frequency/2* cycles per frequency in 4 ms steps for frequencies ranging from 3 to 100 Hz, using ‘*tfr_multitaper’* function from MNE-Python. Time bandwidth for frequency smoothing was set to 2. To receive the desired frequency smoothing the time bandwidth was divided by the time window defined by the number of cycles. For example, for 10 Hz frequency time bandwidth was 2/0.5, resulting in 4 Hz smoothing. We used − 0.3 s to − 0.1 s before the first press as the baseline. The statistical analyses performed on theta (3-7 Hz), alpha (8-14 Hz), β (15-40 Hz), and γ bands (41-100 Hz) used spatiotemporal cluster permutation tests in the same way as for evoked response analyses.

### Cluster based statistical analysis of MEG and EEG data

Cluster-based analyses identified significant clusters of neighboring electrodes or sensors in the millisecond time dimension. To assess the differences between the experimental conditions as defined by behavioral outcomes, we ran cluster-based permutation analysis (Maris & Oostenveld, 2007), as implemented by MNE-Python by drawing 1000 samples for the Monte Carlo approximation and using FieldTrip's default neighbor templates. The randomization method identified the MEG virtual sensors and the EEG electrodes whose statistics exceeded a critical value. Neighboring sensors exceeding the critical value were considered as belonging to a significant cluster. The cluster level statistic was defined as the sum of values of a given statistical test in a given cluster, and was compared to a null distribution created by randomizing the data between conditions across multiple participants. The p-value was estimated based on the proportion of the randomizations exceeding the observed maximum cluster-level test statistic. Only clusters with corrected p-value < 0.05 are reported. For visualization, we have chosen to plot the MEG sensor or the EEG electrode of the significant cluster, with the highest statistical power. For all performed analyses we the same window length (0.4 s), unless stated otherwise in the results section.

### Behavioral data analysis

The analysis of behavioral data was performed using generalized additive mixed models (Wood, 2017; GAMM) as fully described below in the *Single-trial analysis of MEG and EEG data*, unless stated otherwise in the Results section. Each model was fitted with subject as a random factor. For the Block analysis, Block was included as a fixed factor. For the analysis of metacognitive inference, SOJ was entered as a linear predictor of FOJ.

### Binning procedure of behavioral and neuroimaging data

All cluster-based analyses were performed on three conditions defined on the basis of the objective performance in time production (FOJ: short, correct, long) or the subjective self-estimation (SOJ: short, correct, long) separately for each experimental block. Before the binning, the behavioral data were Z-scored on a per block basis to keep the trial count even in each category. Computing these three conditions within a block focused the analysis on local variations of brain activity as a function of objective or subjective performance. To overcome limitations of arbitrary binning, and to capitalize on the continuous performance naturally provided by the time production and the time self-evaluation tasks, we also used a single trial approach, which investigated the interactions between the first and second order terms.

### Single-trial analysis of MEG and EEG data

To analyze single trial data we used generalized additive mixed models (Wood, 2017; GAMM). We briefly introduce the main advantages and overall approach of the method. GAMMs are an extension of the generalized linear regression model in which non-linear terms can be modeled jointly. They are more flexible than simple linear regression models as there is no requirement for a non-linear function to be specified: the specific shape of the non-linear function (*i.e*. smooth) is determined automatically. Specifically, the non-linearities are modeled by so-called basis functions that consist of several low-level functions (linear, quadratic, etc.). We have chosen GAMMs as they can estimate the relationship between multiple predictors and the dependent variable using a non-linear smooth function. The appropriate degrees of freedom and overfitting concerns are addressed through cross-validation procedures. Importantly, interactions between two nonlinear predictors can be modeled as well. In that case, the fitted function takes a form of a plane consisting of two predictors. Mathematically, this is accomplished by modeling tensor product smooths. Here, we used thin plate regression splines as it seemed most appropriate for large data sets and flexible fitting (Wood, 2003). In all presented analyses, we used a maximum likelihood method for smooth parameter optimization (Wood, 2011). GAMM analyses were performed using the *mgcv* R package (Wood, 2009, version 1.8.12). GAMM results were plotted using the *itsadug* R package (Van Rij et al., 2016, version 1.0.1).

Although not widely used, GAMMs are useful for modeling EEG data (Tremblay & Newman, 2015). Here, sensors were not included as fixed effects and the same model was fitted for every sensor separately. The resulting p-values were corrected for multiple comparisons using false discovery rate (FDR) correction (Genovese et al., 2002). For plotting purposes, we averaged the data across significant sensors after FDR correction and refitted the model. The specifics of this refitted model can be found in the tables. Besides typical F and p values, the tables contain the information on the estimated degrees of freedom (edf). Edf values can be interpreted as how much a given variable is smoothed. Although, higher edf values indicate more complex splines, all tested models showed linear splines (edf = 1), depicted in the plotted model outcomes in associated figures.

We fitted the same GAMMs for several neurophysiological measurements chosen on the basis of previous literature. The fitted model contained random effects term for participant and fixed effects that were based on theoretical predictions. Specifically, the full model had the following specification: *uV/T/power* ~ *FOJ* + *SOJ* + *SOJ accuracy* + *FOJ*\**SOJ* + *FOJ*\**SOJ accuracy*. Besides the random term for participants, the model contained smooth terms for the first and second order judgments, *SOJ accuracy* between the first and second order judgment, and the interaction term between *FOJ* and *SOJ accuracy*. Notably, FOJ, SOJ and other predictors were entered as continuous variables in GAM analyses as opposed to post-hoc experimental conditions, which suffered limitations from choosing arbitrary split point in the data.

Although GAMMs have built-in regularization procedures (meaning that they are somewhat inherently resistant against multicollinearity), multicollinearity can been assessed using variance inflation factor (VIF; *fmsb* R package, version 0.5.2). Here, VIF were assessed for the final model and consisted in averaging data from multiple sensors collapsed over a particular variable at hand. None of the VIF values exceeded 1.1, indicating that multicollinearity was unlikely to have had a major influence on the reported findings. Note that Rogerson (2001) recommended maximum VIF value of 5 and the author of *fmsb* recommended value of 10.

Before entering empirical variables in the model, we calculated normalized values or z-scores: trials in which a given variable deviated more than 3 z-scores were removed from further analysis. This normalization was computed separately for every MEG sensor and every EEG electrode. For single-trial analyses of β power in FOJ, we focused on the maximum power within the 0.4 s to 0.8 s period following the R1. This time window overlapped with the selected time window that was used in cluster analyses. For the single-trial analyses of other brain signatures – that is alpha power and sustained activity - we focused on the mean values in time window of 0.4s following or preceding the R2.

## RESULTS

### Participants track the signed magnitude of just produced time intervals

Participants could accurately generate temporal productions (FOJ) with estimates centered around 1.5 s. Fig. 2a provides the normalized (Z-score) FOJ as a function of short, long and correct categories defined according to each individual’s temporal sensitivity (see Methods). To show that participants could accurately self-evaluate their generated durations, we sorted trials on the basis of their self-evaluations (SOJ). We hypothesized that if SOJ and FOJ were independent estimates of endogenous timing, the FOJ sorted as a function of SOJ should differ. Instead, we found the same linear trend when we sorted FOJ on the basis of SOJ (Fig. 2b) as when we sorted times estimates on the basis of FOJ (Fig. 2a). This observation was further corroborated statistically by using a Generalized Additive Mixed Model (GAMM; Wood, 2017) with which we could assess whether SOJ were predictive of FOJ on a single-trial basis. The model fit confirmed that participants could correctly evaluate the signed error magnitude of their generated durations (*F*(4.0) = 192.5, edf = 4.0, p < 10^−15^). These results highlight the main behavioral effect subtending the question of interest in our analyses; complementary behavioral analyses can be found elsewhere (Kononowicz et al., 2017).

### No evidence that ERN is sensitive to the temporal dimension of motor action

The ERN, the seminal electrophysiological signature of error monitoring and self-evaluation, is characterized by a large negative amplitude within 100 ms following an error (Yeung et al., 2004; Holroyd & Coles, 2004). The ERN is obtained by subtracting error trials from correct trials. In the context of our time production task, incorrect trials were the *too short* and *too long* categories. Here, we tested the hypothesis that ERN would reflect a response selection error in the temporal domain (e.g., Luu et al., 2003) and predicted a V-shaped amplitude pattern so that the further away the temporal production was from the target duration, the larger the amplitude of the ERN would be. In short, the larger the error, the larger the ERN amplitude was predicted to be.

For this, we looked at evoked responses immediately following the second keypress (R2), and found a negative deflection peaking at ~60 ms characteristic of an early post-movement activity (Praamstra et al., 2003) which could be seen with both EEG and MEG (Fig. 3 A-B and C-D, respectively). To test the possible sensitivity of ERN to temporal production, we used spatiotemporal cluster permutation tests. We also focused on EEG, which is more sensitive to activity in midline structures such as cingulate cortices, that are the main known sources of the ERN. Contrasts were ran in both neuroimaging modalities for the initial ERN analysis, and source reconstructions combined MEG and EEG signals. We first compared the evoked responses following the production of the time interval (0 to 0.4 s post-R2) as a function of FOJ and SOJ. Statistical tests yielded no significant changes in evoked responses as a function of FOJ or SOJ, whether in EEG (Fig. 3AB; all p > 0.1) or in MEG (Fig. 3CD; all p > 0.1).

**Figure 3.**
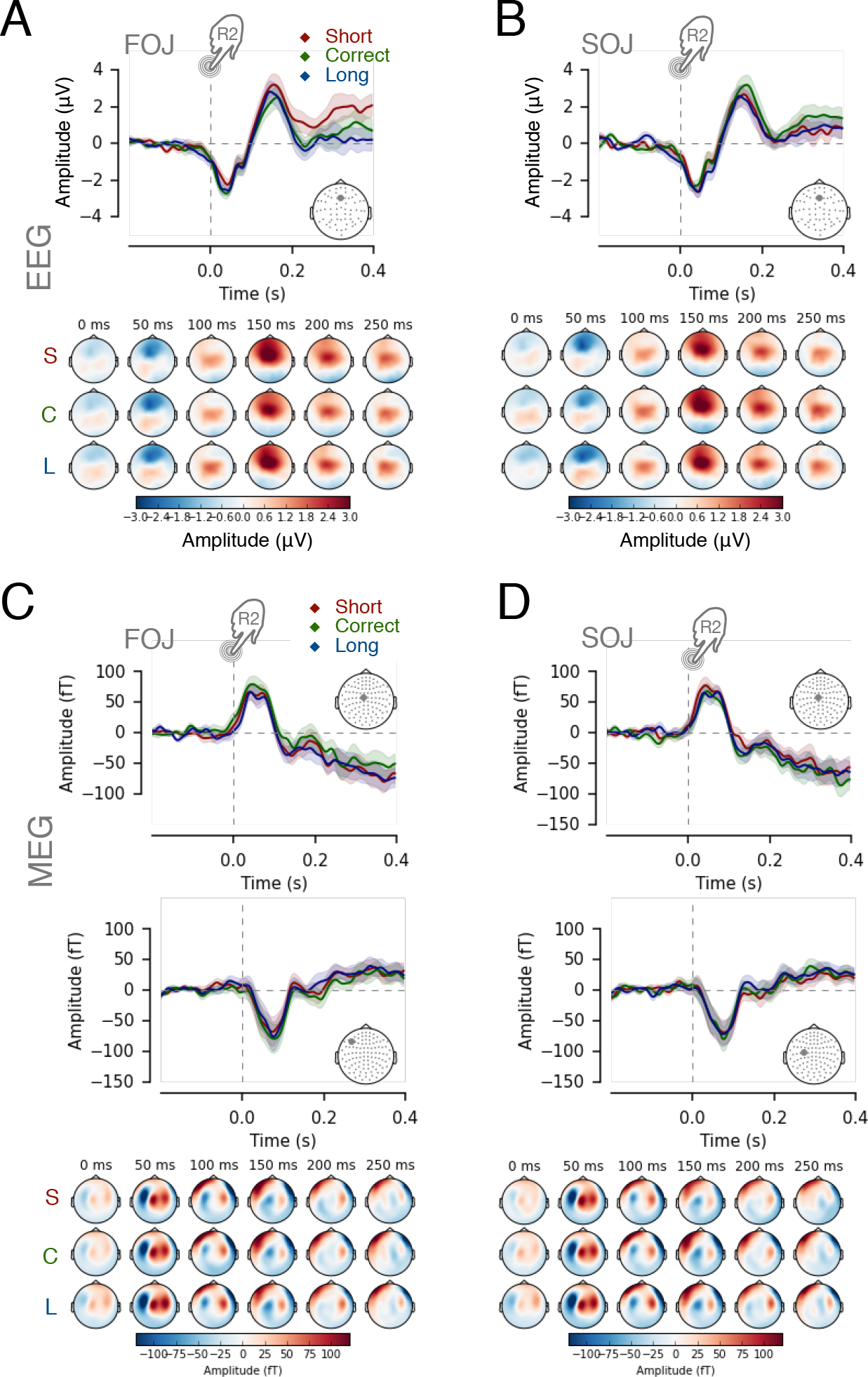
No evidence for ERN-like response. Evoked related potentials / fields (ERP/F) up to 0.4s following R2 showed no significant effect of amplitude as a function of FOJ or SOJ. Time courses plot the gray sensor in the topographical map inset. Colored topographical maps display the data for ‘short’ (S; red), ‘correct’ (C; green) and ‘long’ categories (L; blue). The EEG time course at ‘Fz’ is reported as a function of (**A**) FOJ and (**B**) SOJ categories, respectively. One MEG sensor data is reported as a function of (**C**) FOJ and (**D**) SOJ categories.

Recent work on cognitive control has suggested a link between ERN and theta oscillations (θ: 3-7 Hz) sharing common mid-frontal neural generators (Cohen, 2011; Cavanagh & Franck, 2014). We thus explored evoked θ activity from 0 to 0.2 s post-R2. We constrained the window to 0.2s to prevent capturing spurious activity, extending beyond 0.4 s post-R2, which could originate from the self-evaluation stage. As for the ERN, our prediction was that more theta power would be indicative of larger errors irrespective of their signed magnitude. A cluster permutation test in the θ band (3-7 Hz) yielded no significant changes in θ power as a function of FOJ and SOJ (EEG: p > 0.1 and MEG: p > 0.1). Hence, we found no evidence for a V-shaped pattern as a function of FOJ or SOJ in the ERN or in the θ power that would have supported the TED hypothesis.

### Post-interval oscillatory activity as read-out

While exploring oscillatory responses post-R2, we observed significant clusters in the alpha band power (α; 8-14 Hz) as a function of FOJ categories (Fig. 4A, p = 0.035). The main sources of this effect originated in medial and prefrontal cortices (Fig. 4A, bottom row). We tested the linear relationship between the observed α power and the behavioral variables using a single-trial GAMM analysis on the normalized mean α power. Our analysis revealed a consistent pattern across two distinct clusters, one associated with FOJ, the other with SOJ: the first significant group of electrodes showed a linear relationship between α power and FOJ (F = 22.9, edf = 1, p < 0.0001, Fig. 4B; Table 1) so that shorter trials were associated with a larger α power (*i.e*., the shorter the temporal production, the stronger the α synchronization). In the second significant cluster (Fig. 4c, p = 0.031), trials judged as being ‘too short’ in SOJ were associated with a larger α power (Fig. 4C).

**Figure 4.**
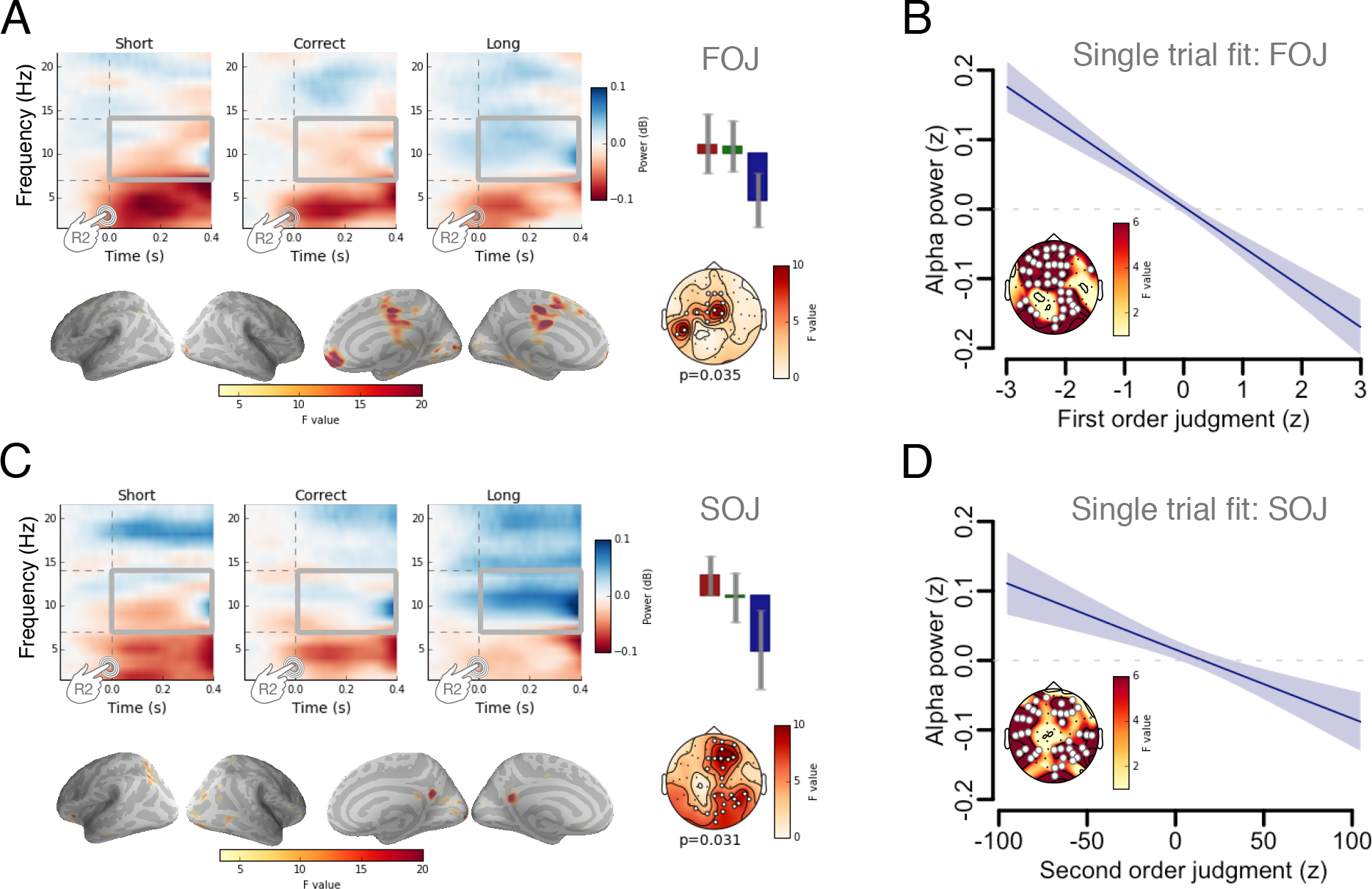
Post-R2 α power signatures of FOJ and SOJ. (**A**) Time-frequency spectra for one EEG sensor illustrating the outcomes of a cluster-based permutation F-test capturing a decrease of α power (8-14 Hz) as a function of FOJ categories. The average α power over the highlighted time window (gray squares) significantly decreased with increasing duration production (bar plot; short: red; green: correct; blue: long). (**B**) The single trial model fit. Post-R2 α power decrease as a function of FOJ. Statistical details in Table 1. (**C**) Time-frequency spectra for the EEG sensor with the largest F-value (cluster-based permutation test). Bar plot illustrating the average α power as a function of SOJ categories: α power decreased with increasing SOJ. Source estimates of the α power effect as a function of SOJ (uncorrected F-map; combined MEG and EEG) implicated medio-central and prefrontal cortices. The strongest source originated from the precuneus. (**D**) The single trial model fit. Post-R2 α power decrease as a function of SOJ. Statistical details in Table 2.

**Table 1.** The results of single trial GAMM analysis based on α power following the second keypress. The table displays the results of the model that was based on the data collapsed across the significant sensors, showing the main effect FOJ, when the model was fitted on per-sensor basis.

| *Parametric coefficients* | *Estimate* | *Std. Error* | *t-value* | *p-value* |
| --- | --- | --- | --- | --- |
| *Intercept* | 0.0042 | 0.0086 | 0.4870 | 0.6262 |
| *Smooth terms* | *edf* | *Ref.df* | *F-value* | *p-value* |
| s(FOJ) | 1.0000 | 1.0001 | 22.9395 | < 0.0001 |
| s(SOJ) | 1.0000 | 1.0001 | 0.3052 | 0.5807 |
| s(SOJ accuracy) | 1.0001 | 1.0002 | 0.0775 | 0.7809 |
| ti(FOJ * SOJ) | 1.0012 | 1.0025 | 0.0019 | 0.9650 |
| ti(FOJ * SOJ accuracy) | 1.0009 | 1.0018 | 3.9561 | 0.0466 |
| s(subject) | 0.0012 | 11.0000 | 0.0000 | 0.9999 |

GAMM analysis: R2-locked $\alpha$ power, FOJ cluster, full model

Importantly, and consistent with the topographical scalp differences, the neural contributors of α power changes in FOJ were distinct from those observed in SOJ (Fig. 4AC): source estimates for the FOJ effect implicated medial, central and prefrontal cortices whereas those for the SOJ effect were located near pre-cuneus. Considering the anatomical separability of the neural generators, we refitted the single trial model without including the FOJ term. This analysis revealed a significant group of electrodes for which α power was linearly predictive of SOJ (F = 12.4, edf = 1, p = 0.0004, Fig. 4D; Table 2).

**Table 2.** The results of single trial GAMM analysis based on α power following the second keypress. The table displays the results of the model that was based on the data collapsed across the significant sensors, showing the main effect SOJ, when the model was fitted on per-sensor basis and the FOJ term has been excluded from the model.

| A. parametric coefficients | Estimate | Std. Error | t-value | p-value |
| --- | --- | --- | --- | --- |
| Intercept | 0.0045 | 0.0080 | 0.5651 | 0.5720 |
| B. smooth terms | edf | Ref.df | F-value | p-value |
| s(SOJ) | 1.0001 | 1.0003 | 12.3827 | 0.0004 |
| s(SOJ accuracy) | 1.0032 | 1.0064 | 0.4280 | 0.5179 |
| ti(FOJ * SOJ) | 1.0016 | 1.0033 | 0.2936 | 0.5888 |
| ti(FOJ * SOJ accuracy) | 2.1676 | 2.6620 | 2.2540 | 0.1042 |
| s(subject) | 0.0012 | 11.0000 | 0.0000 | 0.9994 |

GAMM analysis: R2-locked $\alpha$ power, SOJ cluster, Model without FOJ term

It is noteworthy that both the analysis using categorical responses (*i.e*., data binning as short/correct/long) and the single-trial mixed models indicated that the FOJ and the SOJ effects were topographically distinct. This was also supported by source estimations (Fig. 4C, bottom row). These topographical and source-estimate differences with similar effect size have important consequences for the interpretation of the variations in α power. If the effect in α power shared the same topography for FOJ and SOJ, one interpretation would have been a strong correlation between FOJ and SOJ, which were used as factors in the statistical tests predicting the same variable. Rather, different source estimates and topographical distributions of FOJ and SOJ indicate that distinct brain structures contributed to the evaluation of self-generated behavior.

Hence, following the end of the temporal production (post-R2), changes in α power seemed indicative of the signed magnitude difference between the target interval and the produced time interval. The linear trend of α power following FOJ appeared in line with the *TMC* read-out hypothesis.

### No evidence of access to temporal information before R2

The capacity to report accurately the signed temporal errors, together with the observation of post-R2 α scaling with SOJ and with FOJ, suggested that participants could access their temporal errors. However, a natural question was why, if having access to the signed magnitude of their temporal error, participants did not correct in real-time their temporal production and correct the *when* of their timing generation? Under the *TMC* hypothesis, there would be no *a priori* privileged time at which participants could read-out the internal variable coding for duration, which could be estimated before or after the R2. At ~200 ms preceding a movement (here, R2), recent findings (Schultze-Kraft et al., 2016) have demonstrated that participants could not veto their decision to move. In our experiment, this would signify that information relevant to the outcome of the temporal production task may already be available ~200 ms before R2.

To address whether information relevant to FOJ and SOJ was available for self-evaluation before R2, we assessed R2-locked brain activity (Fig. 5A, evoked and oscillatory from −0.4 to 0 s). The preceding activity in the α band did not show significant clusters, suggesting that information used for self-evaluation was not encoded in the α band before R2.

**Figure 5.**
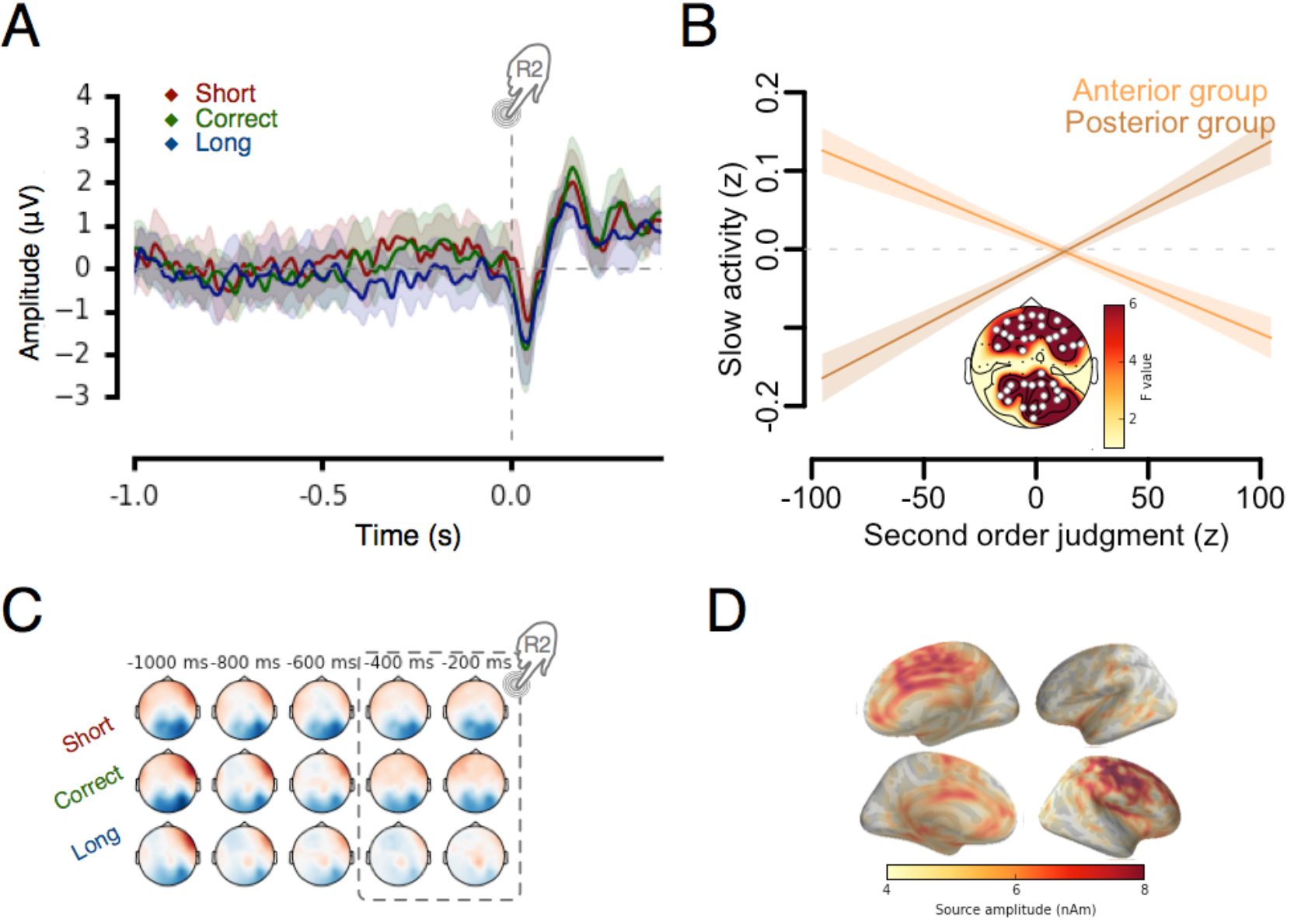
Sustained activity before R2 indicative of SOJ. (**a**) Time courses and topographical plots of EEG activity 1s prior to the second keypress as a function of SOJ. (**b**) Single trial model fit. The sustained brain activity preceding the R2 (−0.4-0s) predict SOJ. The anterior and posterior clusters where plotted separately due to reversed polarities of EEG signals depicted in panel ‘C’. Statistical details in Table 3, 4. (**c**) Topographical maps show significant MEG sensors. (**d**) Cortical source estimates collapsed over all conditions. Motor and mid-frontal regions were the most likely origins of the sustained activity.

At these latencies, on a trial-by-trial basis, we did find significant differences in the amplitude of the evoked activity as a function of SOJ (Fig. 5B): we observed a positive frontal cluster (F = 20.5, edf = 1, p < 0.0001; Fig. 5B, Table 3) that covaried negatively with SOJ (Fig. 5BC) (and a posterior negative cluster that co-varied positively with SOJ; F = 27.7, edf = 1, p < 0.0001; Fig. 5B; Table 4). In line with this bipolar EEG scalp distribution, and in agreement with previous work (Fig. 5C; Wiener *et al.*, 2010), the brain sources at the origin of this activity were located in the motor, premotor, and mid-frontal regions (Fig. 5D).

**Table 3.** The results of single trial GAMM analysis based on sustained activity preceding the second keypress. The table displays the results for the final model that was based on the data collapsed across the significant sensors, showing the main effect SOJ, when the model was fitted on per-sensor basis. The table depicts the anterior cluster.

| *Parametric coefficients* | *Estimate* | *Std. Error* | *t-value* | *p-value* |
| --- | --- | --- | --- | --- |
| *Intercept* | -0.0009 | 0.0072 | -0.1318 | 0.8951 |
| *Smooth terms* | *edf* | *Ref.df* | *F-value* | *p-value* |
| s(FOJ) | 1.0001 | 1.0001 | 0.3260 | 0.5680 |
| s(SOJ) | 1.0002 | 1.0003 | 20.5239 | < 0.0001* |
| s(SOJ accuracy) | 1.0002 | 1.0005 | 1.0091 | 0.3151 |
| ti(SOJ * FOJ) | 1.0027 | 1.0055 | 0.8087 | 0.3700 |
| ti(FOJ * SOJ accuracy) | 1.0089 | 1.0178 | 1.3847 | 0.2368 |
| s(subject) | 0.0015 | 11.0000 | 0.0000 | 0.9948 |

GAMM analysis: R2-locked Readiness potential, Anterior cluster

**Table 4.** The results of single trial GAMM analysis based on sustained activity preceding the second keypress. The table displays the results for the final model that was based on the data collapsed across the significant sensors, showing the main effect SOJ, when the model was fitted on per-sensor basis. The table depicts the posterior cluster.

| *Parametric coefficients* | *Estimate* | *Std. Error* | *t-value* | *p-value* |
| --- | --- | --- | --- | --- |
| (Intercept) | -0.0040 | 0.0074 | -0.5389 | 0.5900 |
| *Smooth terms* | *edf* | *Ref.df* | *F-value* | *p-value* |
| s(FOJ) | 1.0000 | 1.0001 | 0.1091 | 0.7412 |
| s(SOJ) | 1.0000 | 1.0001 | 27.6527 | < 0.0001* |
| s(SOJ accuracy) | 1.0000 | 1.0001 | 4.4611 | 0.0347 |
| ti(SOJ * FOJ) | 2.0365 | 2.5519 | 1.3882 | 0.2115 |
| ti(FOJ * SOJ accuracy) | 1.0077 | 1.0153 | 1.0731 | 0.2974 |
| s(subject) | 0.0003 | 11.0000 | 0.0000 | 0.9815 |

GAMM analysis: R2-locked Readiness potential, Posterior cluster

These analyses suggested that information relevant to subsequent SOJ reports may be elicited and available prior to R2. As the contribution of effective FOJ to slow evoked activity was not relevant, we hypothesize that the sustained activity may reflect an intrinsic decisional biases affecting self-evaluation, functionally distinct from FOJ. To sum up, despite the early formation of SOJ, we do not find convincing evidence to support the notion of the access to temporal information before R2.

### β power timing signature is consistent with post-R2 α scaling

We interpreted the linear scaling between α power post-R2 and SOJ as a strong support of the TMC hypothesis, which predicts that α power should be associated with the internal variable coding for duration. Previous studies have suggested that β power was strongly associated with an internal variable coding for duration (Kononowicz et al., 2017; Wiener et al., 2018). Therefore, the effects observed in the post-R2 activity could be based on the internal variable controlled by β power. To test this hypothesis, we assessed whether β power following the first key press predicted the post-interval α power. This analysis showed that β power was significantly predictive of post-R2 α power in frontal and posterior sensors (F = 44.6, edf = 1, p < 0.0001; Fig. 6; Table 5), strongly suggesting that the value of β fluctuations before at the onset of the temporal production could be read out to elicit the post-interval α modulation (read-out).

**Figure 6.**
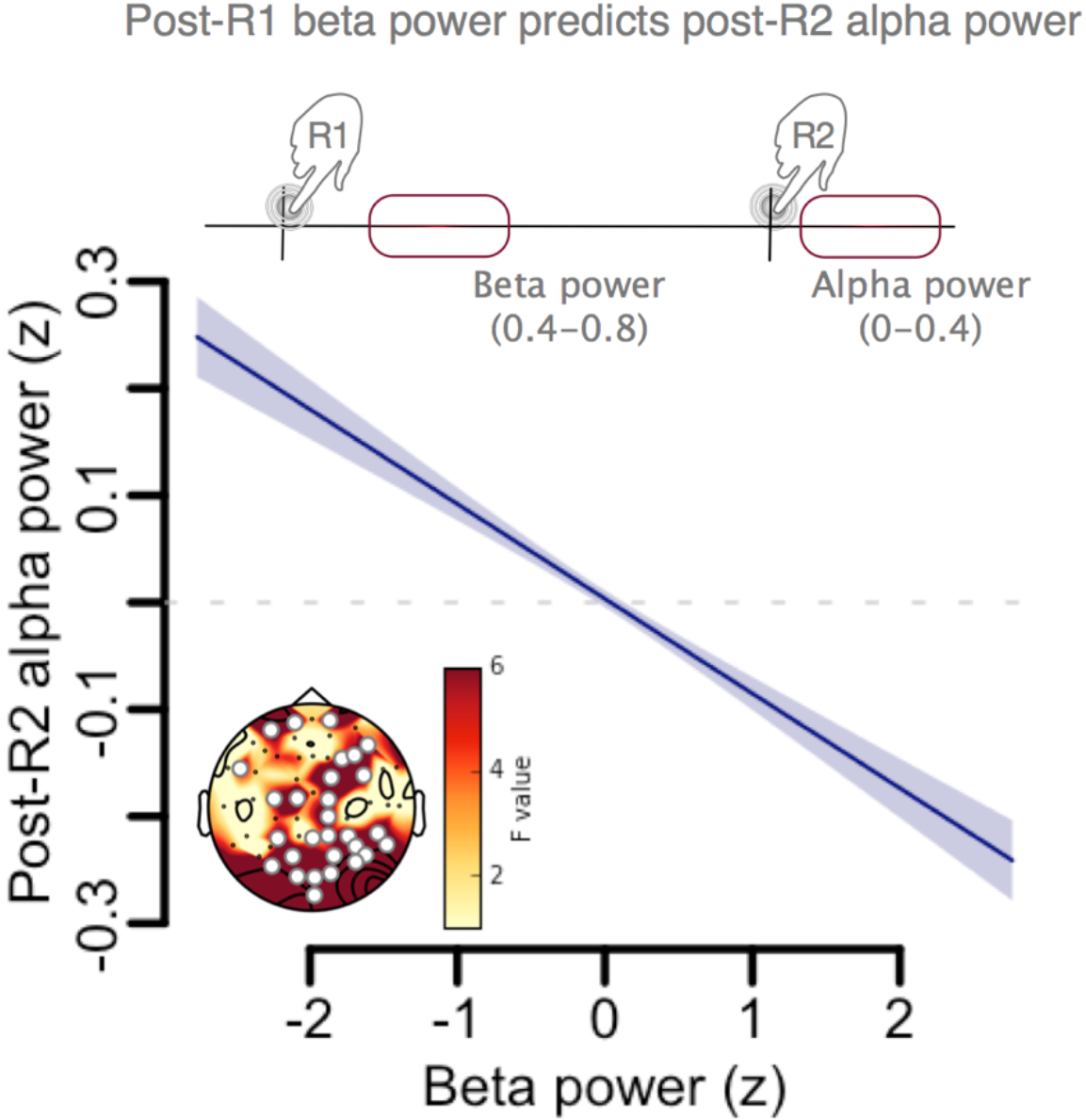
Post-R1 βpower contributes to post-R2 evaluation. Single trial model fit predicting post-Rα2power using the post-Rβ1 significant EEG sensors.

**Table 5.** The results of single trial GAMM analysis where beta power was tested as a predictor for post-R2 alpha. The table displays the results for the final model that was based on the data collapsed across the significant sensors.

| *Parametric coefficients* | *Estimate* | *Std. Error* | *t-value* | *p-value* |
| --- | --- | --- | --- | --- |
| (Intercept) | 0.0034 | 0.0079 | 0.4275 | 0.6691 |
| *Smooth terms* | *edf* | *Ref.df* | *F-value* | *p-value* |
| s(Beta power) | 1.0002 | 1.0004 | 44.5739 | < 0.0001 |
| s(subject) | 0.0032 | 11.0000 | 0.0000 | 1.0000 |

GAMM analysis: Beta predicting post-R2 alpha

In sum, we identified distinct cortical signatures of self-evaluation of temporal production. Moreover, post-interval α desynchronization was linked to preceding β activity, suggesting an evaluation of internal variable coding for duration as previously suggested by the accurate representation of individuals’ temporal uncertainties (Balci et al., 2009).

## DISCUSSION

We assessed two working hypotheses on the mechanisms supporting the evaluation of self-generated time intervals (TED and TMC), using a task in which participants produced durations, and self-evaluated the signed error magnitude of their estimates while being recorded with combined MEG and EEG. We found no supporting evidence for the generation of an ERN modulated as a function of temporal error in this task; however, we found that α power following R2 negatively correlated with SOJ and with FOJ. We interpret these findings as evidence in favor of the TMC working hypothesis: the neuronal signature following R2 may linearly correlate with the signed difference between the internal variable coding for duration, and the target duration. In support of the TMC hypothesis, the initial β power, known to scale with duration of a produced interval in this task (Kononowicz et al., 2017), predicted the post-R2 α power. Below, we discuss these interpretations together with the current shortcomings of our study.

### Temporal meta-representations

Temporal metacognition posits the existence of a process that actively infers the state of an internal variable coding for a target duration, *i.e*., the meta-representation of a duration (van Wassenhove, 2009). This meta-representation entails the extraction of a first order variable from the initial state of the system (Cleeremans et al., 2007). What crucially follows is that the meta-representation of duration would be specified by neural signatures that would be anatomically and functionally distinct from the neural signatures of the first order variable (Lak et al., 2014).

In line with this assertion, the anatomical separation of the post-R2 α power generators and the post-R1 β power generators (Kononowicz et al., 2017) was visible in source estimates, fulfilling the criterion of anatomical separability between the first and second order representations of duration. Second, post-R2 α power and post-R1 β power operated at different latencies, further strengthening the dissociation of first and second order representations along the temporal dimension. Thirdly, although both signatures of inhibition, α and β neural oscillations are typically ascribed different functional roles, with α implicated in global network regulation (Palva & Palva, 2012), and β implicated in the representation of various sensorimotor features (Engel & Fries, 2010; Kilavik et al., 2013). Together, these features of post-R2 α and post-R1 β suggest the possibility of a meta-representation of duration.

Indirect evidence also suggests the existence of temporal meta-representations, that may require a read-out process with the read-out understood as being either active or passive (Fleming & Daw, 2017): a passive sensitivity to the state of the system would be in line with the TED hypothesis, whereas TMC would predict an active process. Essentially, the post-R2 α decrease may reflect the outcome of an active read-out process. One limitation of the present study is that there were no trials in which participants did not self-evaluate their time production. If the metacognitive read-out is an active process, the absence of self-evaluation should abolish the post-R2 α as a function of duration category. Fortunately, previous EEG work assessing time reproduction tasks in the absence of metacognitive inference (Kononowicz & Van Rijn; 2015; Fig. 4 and 5, lower panel) showed no post-R2 alpha or theta power changes as a function of duration category. Thus, changes in post-R2 α may only be seen when participant are explicitly asked to self-evaluate their temporal performance which would be in line with an active metacognitive read-out.

Additional evidence further support the idea that first order signals could be read-out by second order areas. For example, pulvinar neurons have been shown to encode confidence, a second order variable, independently of other areas processing first order variables (Komura et al., 2013): this study suggested that one population of neurons can read-out the activity of neural population encoding primary sensory variables. Other studies have also suggested that particular brain regions independently code for first and second order signals (Lak et al., 2014). In humans, similar notions have been explored: using TMS, prefrontal areas have been shown to read-out the strength of perceptual signals in service of confidence judgments (Shekhar & Rahnev, 2018). In line with these ideas, the mapping between β power and duration may be realized via networking through higher order brain regions. For example, prefrontal cortex, implicated in timing (Kim et al., 2017), could monitor signals in motor cortex (Narayanan & Laubach; 2006) or the cortico-basal ganglia loop. Indeed, the cortical sources observed in our study were consistent with the acknowledged role of midline cingulate regions in self-monitoring (Miyamoto et al., 2017), and error monitoring (Ullsperger et al., 2014). Moreover, the orbitofrontal and posterior cingulate were implicated in the metacognitive performance: the association between FOJ and α power originated from prefrontal cortices and ACC, whereas the association between SOJ and α power implicated the precuneus, which has been reported during confidence judgments (De Martino et al., 2013; Ye et al., 2018), and error processing (Menon et al., 2001). Interestingly, gray matter volume in precuneus predicts introspective accuracy (Fleming et al., 2010), and metacognitive efficiency in memory (McCurdy et al., 2013).

### What signals could be read out

Previous studies reported that β power scaled with self-generated durations (Kononowicz et al, 2017; Kononowicz & Van Rijn, 2015), and that the degree of separation in β power predicted individuals’ temporal metacognition performance (Kononowicz et al., 2017). As β power carries internal duration signals, we hypothesized that it could be a signature that the read-out process could rely on and we found that β power predicted the post-R2 α power.

Finding the strongest effects in α power suggests that the monitoring of internal states could rely on different sources of information than just β power, and two studies further support this notion. First, α oscillations have been implicated in performance monitoring when task errors relied more on the attentional lapses than on the lack of executive control over the motor system (van Driel et al., 2012). Second, participants can monitor their attentional state, *i.e*. have an appropriate metacognition of an internal variable, which depended on the lateralization pattern of α oscillations (Whitmarsh et al., 2014). According to this view, the post-R2 α decrease observed here could be seen as a reorienting of attention following the generation of durations that were too long. Specifically, in an attentional gate model of time perception (Zakay & Block, 1995), long durations would correspond to not enough attention paid to temporal production, hence more attention needed for the next trial. However, the lack of association between α power during the self-generation of durations and produced duration in this dataset (Kononowicz *et al.*, 2017) and in previous studies (Kononowicz & Van Rijn, 2015) do not fit well with this interpretation. Whether α power could contribute to metacognitive performance in the timing of longer durations thus remains to be tested.

Another open question is why read-out related processes and self-monitoring would implicate α oscillations. When assessing the role of cross-frequency coupling in this timing task, we found that the coupling strength between the phase of alpha oscillations and the power of beta oscillations, was indicative of the precision with which participants self-generated a duration (Grabot, Kononowicz, et al, 2017). This pattern was found during the generation of the interval. One speculative hypothesis is that the termination of the interval (R2) may implicate the read-out of the precision maintained in the coupling of β power with respect to the phase of alpha. This mechanism would be close to the prediction of oscillatory-based mechanisms in event timing which implicate the phase of oscillations in timing precision (Gallistel, 1990). This, however, remains an open and very difficult question to address empirically, one for which a set of dedicated experiments – including animal work – would be needed.

### Why participants do not correct temporal errors if they have access to it?

If access to temporal error is possible early on in the trial, why didn’t participants correct for their ongoing interval production before terminating their interval production? We found changes in slow evoked activity locked to R2 that scaled with the metacognitive judgment, and considered that it presumably initiated the read-out process. However, this information did not appear relevant to the decision yielding to the termination of R2. Together with the post-R2 α power scaling with FOJ and SOJ, these results suggest that participants may only have access to their temporal errors after the time production termination. However, partial correction may still be a viable possibility, in which case such corrective behavior would be of great importance to timing models and additional studies will address this question

### Conclusions

In summary, our results provide novel insights with the possibility that meta-representations of duration estimates are viable to support of temporal metacognition during self-generated duration production. In line with the prediction of temporal metacognition (TMC hypothesis), we found a linear scaling between produced duration and the neural responses elicited after the temporal production (post-R2), suggesting neuronal signatures of read-out of internal variable coding for duration.

## AUTHOR CONTRIBUTIONS

V.W., T.W.K. designed the research and performed the experiments. T.W.K. and V.W. analyzed the data. T.W.K. and V.W. wrote the manuscript.

## ACKNOWLEDGMENTS

This work was supported by an ERC-YStG-263584 and an ANR10JCJC-1904 to V.vW. We thank the members of UNIACT and the medical staff at NeuroSpin for their help in recruiting and scheduling participants. We thank Clémence Roger for her initial contributions to the study, members of UNICOG for fruitful discussions. Preliminary results were presented at SFN (2016).

